# Characterization of Mucosal Dysbiosis of Early Colonic Neoplasia

**DOI:** 10.1101/662221

**Authors:** Bo-young Hong, Takayasu Ideta, Yuichi Igarashi, Yuliana Tan, Michael DiSiena, Allen Mo, John W. Birk, Faripour Forouhar, Thomas J. Devers, George M. Weinstock, Daniel W. Rosenberg

## Abstract

Aberrant crypt foci (ACF) are the earliest morphologically identifiable lesion in the colon that can be detected by high-definition chromoendoscopy with contrast dye-spray. Although frequently associated with synchronous adenomas, their role in colorectal tumor development, particularly in the proximal colon, is still not clear. The goal of this study was to evaluate the profile of colon-associated bacteria associated with proximal ACF and to investigate their relationship to the presence and subtype of synchronous polyps present throughout the colon. Forty-five subjects undergoing a screening or surveillance colonoscopy were included in this retrospective study. Our study cohort included a total of 16 subjects with no identifiable proximal lesions (either ACF or polyp), 14 subjects with at least 1 ACF but no polyp(s), and 15 subjects with both at least 1 ACF and a synchronous proximal polyp(s) detected at colonoscopy. Bacterial cells adherent to the epithelia of ACF and normal mucosa were visualized by *in situ* hybridization within confocal sections. Bacterial DNA isolated from biopsies was used to construct PCR amplicon libraries targeting the V4 hypervariable region of the 16S rRNA gene, which were then sequenced on the Illumina platform. ACF showed significantly greater heterogeneity in their bacterial profiles compared to normal mucosa. Interestingly, one of the bacterial community structures we characterized was strongly correlated with the presence of synchronous polyps. The observed dysbiosis is more prominent within the colonic epithelium that also harbors synchronous polyps. Finally using DNA-mass spectrometry to evaluate a panel of colorectal cancer hot-spot mutations present in the ACF, we found that several *APC* gene mutations (*R1450*, R876*, S1465fs\**3) were positively associated with the presence of *Instestinibacter sp*., whereas *KRAS* mutations (G12V, G12D) were positively correlated with *Ruminococcus gnavus*. This result indicates a potential relationship between specific colon-associated bacterial species and somatically acquired CRC-related mutations. Overall, our findings suggest that perturbations to the normal adherent mucosal flora may constitute a risk factor for early neoplasia, demonstrating the potential impact of mucosal dysbiosis on the tissue microenvironment and behavior of ACF that may facilitate (or impede) their progression towards more advanced forms of neoplasia.

## Background

Colorectal cancer (CRC) is the third-leading cause of cancer-related deaths in the US. Fortunately, the widespread application of screening colonoscopy, together with the identification and removal of precancerous polyps, has led to a significant overall reduction in CRC incidence ^1–3^. Despite the overall health benefits, however, endoscopic surveillance has failed to uniformly prevent the occurrence of CRC, particularly in the proximal colon ^4^. These limitations are underscored by patients who develop "interval" CRC between screening colonoscopies and subsequent surveillance. Most of these cases have been attributed to non-detected or incompletely resected proximal colon lesions that were initially present during the index colonoscopy ^5–8^. Thus, there is a need to develop more robust strategies that enable the accurate identification and removal of proximal colon lesions and importantly, to enable the identification of those individuals at increased risk for recurrent neoplasms at the time of index colonoscopy ^9^.

Aberrant crypt foci (ACF) are the earliest morphologically detectable lesion that are frequently present in the colon ^9^. However, ACF are not routinely detected during conventional colonoscopy due to their diminutive size (<5 mm in diameter) ^10–14^. Despite evidence that ACF are often associated with the presence of adenomas, their role in tumor development, particularly in the proximal colon, is still actively debated ^15–18^. Our laboratory has recently demonstrated the power of high-definition chromoendoscopy to identify colonic ACF within the proximal colon ^19^. We further validated an ultra-sensitive DNA-mass spectrometry platform that, combined with laser-capture microdissection (LCM), enables the detection of somatic mutations across a wide panel of CRC-related oncogenes and tumor suppressor genes ^9,19^. We believe these recent studies, including our genome-wide methylation analysis of ACF ^20^, firmly establish the premalignant potential of ACF and their potential application to risk prediction.

During the past two decades, there has been a growing appreciation for the role of the gut microbiome in CRC pathogenesis ^21^. *Fusobacterium nucleatum* was identified as an abundant taxon from colorectal tumor tissues, while other species were also detected ^22,23^. *Escherichia coli* has also been associated with CRC. Colonic biopsies of adenomas and CRCs showed the increased presence of intracellular *E. coli* ^24^, while higher abundance of other bacterial species of the phylum Proteobacteria were found in rectal adenomas ^25^. Enterohemorrhagic *E. coli* and enteropathogenic *E. coli* are also known to be risk factors for CRC, most likely by generating toxins that affect colonic tissue ^26,27^. Toxin-producing *Bacteroides fragilis* can impact colorectal carcinogenesis by its production of *B. fragilis* toxin, resulting in disruption of E-cadherin junctions, β-catenin signaling and IL-8 expression. ^28,29^.

Our understanding of the characteristics of the colon-associated microbiome and its potential role in the development of early colonic neoplasia remain incomplete. The majority of existing studies have examined microbiota in stool samples and/or focused on a limited set of bacterial taxa. However, we believe this approach may have limited the amount of information that can be obtained by directly analyzing the composition of the colon-associated bacterial community present on mucosal lesions during early neoplasia. Since mucosal adherent bacteria are likely to play a more direct role in the pathogenesis of CRC than luminal bacteria ^30^, robust approaches that take into account potential interactions with distinct bacterial taxa and their predicted metabolites will be required for a broader understanding of the complex etiology and progression of this disease. Our long-term goal is to determine whether the microbiome can influence the microenvironment and behavior of early colonic neoplasia, either facilitating (or impeding) the progression of small lesions to more advanced forms of neoplasia.

## Methods

### Study population and endoscopic procedure

Eligible healthy adults (50 to 65 years of age) that were referred to the Division of Gastroenterology at the John Dempsey Hospital/University of Connecticut Health (Farmington, CT) for screening or surveillance colonoscopy were recruited to the ACF study by participating physicians during their initial office consultation. Patients were screened for a standard-of-care colonoscopy procedure. Mucosal biopsy samples were procured under strict guidelines approved by the Institutional Review Board from the University of Connecticut Health (UCH) Committee (#IE-10-068SJ-3.2) between 2010-2017. All participants provided written informed consent prior to inclusion in the study. Patients who met the Amsterdam criteria for familial adenomatous polyposis (FAP) or hereditary non-polyposis CRC (HNPCC) were excluded. Patients with ulcerative colitis, active infectious gastroenteritis, proctitis, diseases of malabsorption or a history of CRC were excluded. To limit the confounders of age and smoking, all subjects selected were non-smokers. Prior to colonoscopy, all participants completed a study questionnaire, including information on smoking, current medications and supplement use, previous history of endoscopy, and family history of cancer. Body mass index (BMI, kg/m^2^) was calculated from weight (kg) and height (m) measurements obtained during the initial office consultation.

High-definition chromoendoscopy was performed in the distal 20-cm of the colorectum and throughout the entire proximal colon with a freshly prepared solution of 0.1% indigo carmine dye-spray, using a spray catheter for contrast enhancement. The identification and histologic evaluation of ACF has been described in our previous publications ^9,31^. ACF were isolated from grossly normal-appearing colonic mucosa by biopsy *in situ* and removed using biopsy forceps. In addition, each subject had a histologically confirmed corresponding normal biopsy specimen taken from the same region of the colon, generally within 2-cm of the ACF biopsy sample. ACF were identified if two or more crypts had an increased luminal diameter of 1.5 to 2 times the luminal size of surrounding crypts. ACF were visualized, videotaped and photographed using an Olympus high-definition colonoscope. Biopsies of individual ACF and/or normal colonic mucosa were embedded immediately in optimal cutting temperature (OCT) media and stored at −80°C until further analysis. Samples were stained with hematoxylin and eosin (H&E) and ACF were histologically confirmed by a board-certified GI pathologist (F.F.).

### Confocal microscopy

Human tissues were stained by fluorescence *in situ* hybridization using a 16S rRNA probe as previously described (28). Briefly, OCT-embedded tissue was cryo-sectioned at a 5-μm thickness and then fixed in 4% paraformaldehyde for 24 hours at 4°C. Formalin-fixed paraffin-embedded (FFPE) human polyp tissues were de-paraffinized with sequential washes in xylene plus ethanol. Samples were then hybridized using the EUB338-cy3 probe (Eurofins Genomics) at 55°C overnight. Immunofluorescence was performed with 5% BSA blocking, followed by staining with anti-Mucin-2 (Santa Cruz Biotechnology) and anti-E-cadherin (Cell Signaling Technology) at 4°C overnight. Alexa Fluor 488 (Thermo Fisher) and Alexa Fluor 647 (Thermo Fisher) were used to detect Mucin-2 and E-cadherin, respectively. Nuclei were stained with 10μg/mL DAPI (Sigma-Aldrich). Slides were mounted with Prolong Gold (Invitrogen). Images were taken at 63x/1.4A magnification using a ZEISS LSM 880 Confocal Microscope maintained at the CCAM Microscopy Facility at the University of Connecticut Health Center. Images were captured with z-stack and 10% overlap tile-scanning to obtain in-depth bacterial localization on each mucosal sample. Finally, images were processed with ImageJ to generate a final image.

### DNA extraction, 16S rRNA gene amplicon library construction and sequencing

DNA was extracted from whole tissue biopsies using the MoBio PowerMag Soil 96 well kit (MoBio Laboratories) according to the Manufacturer’s protocol for the Eppendorf epMotion liquid handling robot. DNA extracts were quantified using the Quant-iT PicoGreen kit (ThermoFisher Scientific). The V4 hypervariable region of the bacterial 16S rRNA gene was amplified using 30 ng of extracted DNA as a template and the primer set of 515F and 806R with Golay code indices ^32^. Samples were amplified in triplicate using GoTaq (Promega) with the addition of 10 μg of BSA (New England BioLabs). The PCR reaction was incubated at 95°C for 3.5 minutes with 30 cycles of 30 s at 95.0°C, 30 s at 50.0°C and 90 s at 72.0°C, followed by a final extension at 72.0°C for 10 minutes. PCR products were pooled for quantification and visualization using the QIAxcel DNA Fast Analysis (Qiagen). PCR products were normalized based on the concentration of DNA from 250 to 400-bp, then pooled using the QIAgility liquid handling robot. The pooled PCR products were cleaned-up using the Mag-Bind RxnPure Plus (Omega Bio-tek) according to the Manufacturer’s protocol. The cleaned pool was sequenced on the MiSeq (Illumina, Inc.) using the v2 2×250 base-pair kit (Illumina, Inc.). Positive/negative controls were included for DNA extraction and amplification.

### Sequence data processing

Raw 16S rRNA gene reads were processed in mothur ^33^. Sequencing reads were processed by removing the sequences with low quality (average quality <25) and ambiguous bases (N’s). Chimeric amplicons were removed using UChime software (https://omictools.com/uchime-tool). For 16S, an operational taxonomic unit (OTU)-based approach was used by clustering sequences with 3% dissimilarity cutoff in Usearch ^34^ and the taxonomic classification *via* the Ribosomal Database Project Classifier ^35^. Shannon diversity index was calculated for each sample and visualized using GraphPad Prism software. Significant separation of microbiome samples based on pair-wise thetaYC distance was calculated, visualized as principal coordinates analysis (PCoA) plots using R (https://www.r-project.org/). Hierarchical clustering of ACF and normal mucosal bacterial communities based on OTU relative abundances was performed *via* Jaccard distances and the average linkage method in Morpheus (https://software.broadinstitute.org/morpheus). RAWGraph was utilized to visualize microbiome clusters A and B.

### Predictive functional profiling

Predictive functional profiling was performed using a phylogenetic investigation of communities by reconstruction of unobserved states (PICRUSt) ^36^ inferred from 16S rRNA gene copies as OTUs. Predicted functional gene contents or Kyoto Encyclopedia of Gene and Genomes (KEGG) Orthologs (KO) of bacterial communities were estimated further to predicted functional pathways using *“predict_metagenomes.py”* command of PICRUSt with default parameters. Differentially featured gene function was selected using LEfSe ^37^ with the default settings. Results were filtered to contain a logarithmic LDA score > 2.5.

### Mutation analysis

The customized DNA-mass spectrometry assay design included the following CRC hot-spot somatic mutations; *APC_R1450X, APC_S1465fsX3, APC_R876X, KRAS_G12DV, NRAS_G12D, NRAS_G13D, BRAF_V600E, ERBB2_G776insYVMA-c2325* and *ERBB2_G776insYVMA-c2324* ^9^. Mutation screening was performed on a fee-for-service basis by the Genomics Shared Resources Core facility at the Roswell Park Comprehensive Cancer Center ^38^. Briefly, the protocol involves PCR amplification of DNA using SNP-specific primers, followed by a base-extension reaction using the iPLEX PRO chemistry (Agena Bioscience). The PCR products were treated with (shrimp alkaline phosphatase (SAP), then temperature-ramped to 85°C for 5 min to remove excess dNTPs. iPLEX PRO extension enzyme (Agena) was used for the base-extension reactions. The final base-extension products were treated with SpectroCLEAN (Agena) resin to remove contaminating salts. The extension product was spotted on a 384-pad SpectroCHIP II (Agena) using a Sequenom MassARRAY Nanodispenser (Agena, San Diego, CA). A MassARRAY Analyzer Compact MALDI-TOF MS (Agena) was used for data acquisition. All resultant genotyping calls were performed by the MassARRAY Typer Analyzer v4.0.26.73 (Agena).

### Statistical analyses

The significance for PCoA plot clustering was tested *via* analysis of molecular variance (AMOVA)^39^. Differentially abundant taxa between ACF and normal mucosa within subject, across subject, and based upon mutation subtypes were identified *via* Wilcoxon Signed Rank test or the Mann-Whitney test as appropriate. All statistical tests were adjusted for multiple comparisons using the Benjamini-Hochberg false discovery rate (FDR) method. Correlations between the sum of relative abundances of microbiome clusters A or B signature OTUs and clinical parameters were performed *via* Spearman rank-order tests.

## Availability of data and material

Sequence data that support the findings of this study have been deposited in SRA (PRJNA511474).

## Results

### Description of clinical and demographic information

For this study, we retrospectively selected a total of 45 patients who had undergone a routine screening colonoscopy at the JDH. We assigned the patient samples to the following three experimental groups: Group I (n=16 patients) had no identifiable lesions (ACF or polyps) present in the proximal colon; Group II (n=14) had at least one proximal ACF detected at colonoscopy, but no synchronous polyps; Group III (n=15) had at least one or more proximal ACF and synchronous proximal polyp(s) detected at the time of colonoscopy. The characteristics of the study population are shown in **Table 1**.

**Table 1.** Clinical and demographic characteristic of subjects included in the study.

|  | **Normal** |  | **ACF only** |  | **ACF + Polyp** |  |
| --- | --- | --- | --- | --- | --- | --- |
| **Sample size (%)** | n= 16 | (%) | n= 14 | (%) | n= 15 | (%) |
| **Age** | 55.9±9.9 |  | 55.9±7.5 |  | 57.5±7.7 |  |
| **Male** | 8 | (50.0) | 3 | (21.4) | 8 | (53.3) |
| **BMI (kg/m<sup>2</sup>)** | 30.0±7.0 |  | 29.5±6.7 |  | 30.9±6.8 |  |
| **Caucasian** | 9 | (56.3) | 10 | (71.4) | 14 | (93.3) |
| **African American** | 7 | (43.8) | 4 | (28.6) | 1 | (6.7) |
| **Smoking History** | 9 | (56.3) | 6 | (42.9) | 9 | (60.0) |
| **Current Smoker** | 3 | (18.8) | 2 | (14.3) | 3 | (20.0) |
| **Family History of Colon Cancer** | 5 | (31.3) | 6 | (42.9) | 3 | (20.0) |
| **Diabetes** | 4 | (25.0) | 3 | (21.4) | 2 | (13.3) |
| **Daily Regular Strength Aspirin (325 mg) use** | 5 | (31.3) | 3 | (21.4) | 5 | (33.3) |
| **Daily baby Aspirin (81 mg) use** | 6 | (37.5) | 4 | (28.6) | 6 | (40.0) |
| **Regular other NSAIDs use** | 5 | (31.3) | 8 | (57.1) | 7 | (46.7) |
| **Regular multivitamin use** | 9 | (56.3) | 9 | (64.3) | 4 | (26.7) |
| **Regular folic acid use** | 3 | (18.8) | 1 | (7.1) | 1 | (6.7) |
| **Regular calcium use** | 3 | (18.8) | 6 | (42.9) | 3 | (20.0) |
| **Regular vitamin D use** | 3 | (18.8) | 1 | (42.9) | 7 | (46.7) |
| **Antibiotics use (within 1 month)** | 1 | (6.3) | 1 | (7.1) | 0 | (0.0) |

NOTE: Values are means ± SD or n, %.

### Direct visualization of colon-associated bacteria within the colonic mucosa

ACF, adjacent normal mucosa and synchronous polyps in Group III were directly examined for the presence of adherent bacteria using 16S universal FISH probes. Representative lesions are shown in **Figure 1**(ACF and adjacent normal) and also in **Supplemental Figure 1** (synchronous polyps identified in Group III). Bacteria were observed within the mucous layer (green; mucin-2 positive) of the colonic epithelial lining in normal, ACF and polyp tissues. However, in no cases did we observe bacterial infiltration into the colonic epithelial lining nor within the stroma (**Figure 1 and Supplemental Figure 1**). In addition, colon-associated bacteria showed no detectable differences morphologically.

**Figure 1.**
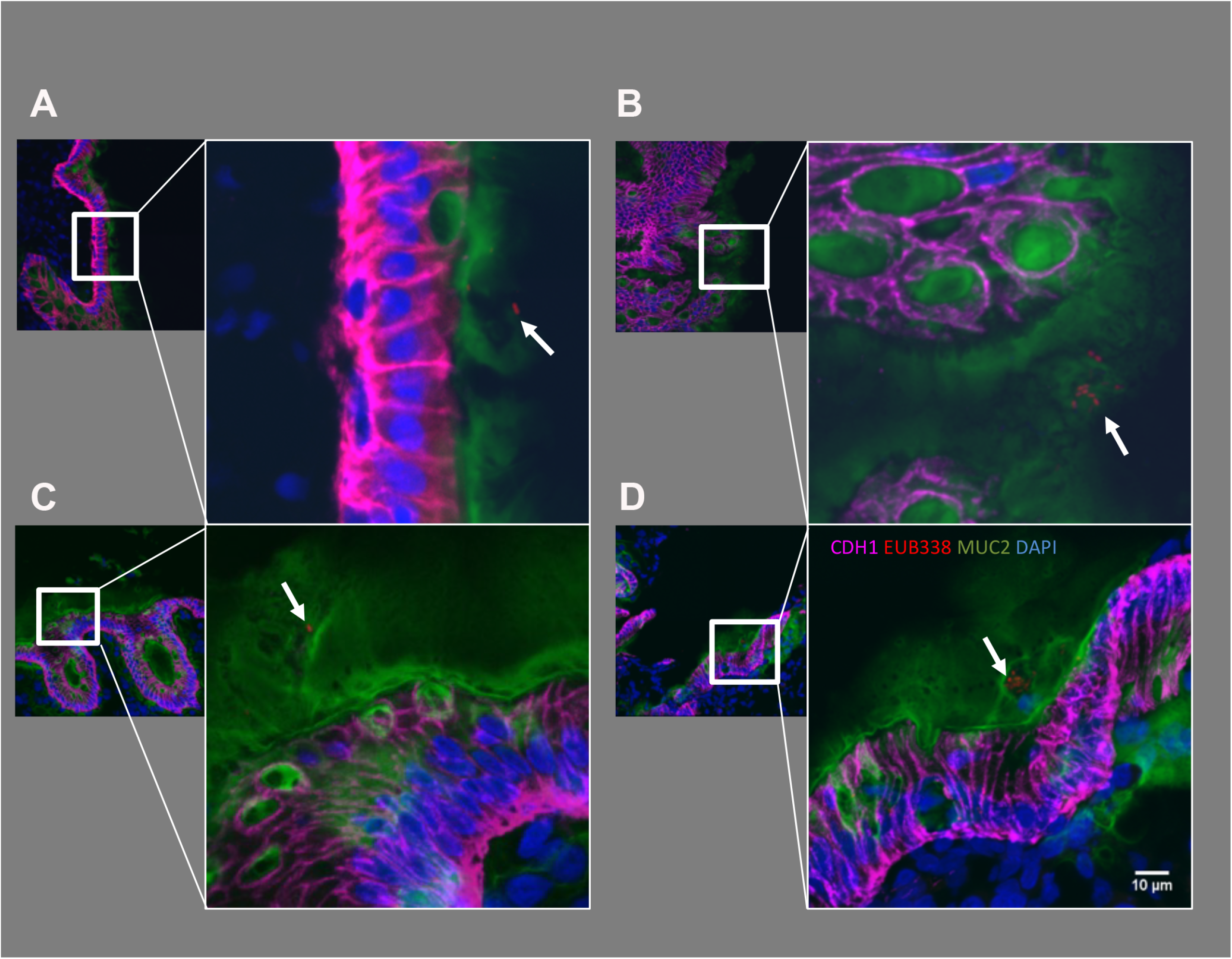
Direct visualization of tightly colon-associated bacteria within the colonic mucosa. ACF and adjacent normal mucosa were directly examined for the presence of colon-associated bacteria using 16S universal FISH probes. Clusters of bacteria were observed within the mucous layer on epithelium. Red: bacteria (EUB338-cy3 probe), Magenta: E-cadherin, Green; Mucin-2, Blue: DAPI. Bacteria are marked with white arrows. Bacteria were observed within the mucous layer associated with the epithelium.

### Between-group analysis of proximal ACF and colon-associated microbiome

Illumina sequencing of the V4 hypervariable region of 16S rRNA amplicons from all individual samples yielded 7,469,049 raw reads and 4,414,345 reads after pre-processing of the data-set. The sequence counts per sample ranged from ~5,627 to 319,577 reads. Libraries were normalized by random sub-sampling for comparisons across the samples. In total, 815 OTUs were found at a 97% identity cut-off from 74 samples. In order to determine alpha and beta diversity, we compared the adherent bacteria of ACF to adjacent normal colonic mucosa from patients with or without polyps, separately. We also compared adherent bacteria of ACF to the normal colonic mucosa of 16 control patients who had no detectable proximal ACF.

First, alpha-diversity analysis (**Figure S2A**) showed median richness of 100.5 and 103.5 different taxa present in the ACF lesion from patients without synchronous polyps and their paired adjacent normal mucosa, respectively. ACF lesions and adjacent normal mucosa from patients with synchronous polyp(s) showed median richness of 103.3 and 99.6 distinct bacterial taxa, respectively. These results were slightly lower than median richness of 110 bacterial taxa found in normal mucosa from the lesion-free patients. However, there was no statistically significant difference in bacterial richness between these groups. In addition, diversity and evenness of the bacterial community within each sample showed no statistically significant differences (**Figure S2B-C**). Nonetheless, subject variability of the bacterial microbiome in lesions from patients with a polyp was consistently greater than lesion-free patients for richness, diversity and evenness.

Next, we characterized the bacterial community structure of ACF biopsies taken from the proximal colon with or without synchronous polyps (**Figure 2**). Pair-wise comparison of bacterial community structure based on the thetaYC index, which takes into account membership of taxa and their relative abundance, was measured between samples. Bacterial community structure visualized in PCoA plots showed no distinct clustering of ACF sites compared to their normal adjacent mucosa or normal mucosa from ACF lesion-free patients (p= 0.139) (**Figure 2A**). However, there was a trend observed in which control mucosal samples (blue) clustered more tightly than ACF samples (red) or their adjacent normal mucosa (yellow), suggesting that normal mucosa is somewhat more homogeneous than the ACF lesions in terms of bacterial diversity (**Figure 2A**). Furthermore, we confirmed that the distance measured from pair-wise thetaYC dissimilarity comparisons between every sample in both ACF sites (purple) and their adjacent normal mucosa (green) were significantly different from that of control normal mucosa (blue) (p< 0.0001, p= 0.008) (**Figure 2B**), indicating that ACF patients harbor a significantly more heterogeneous bacterial structure than control subjects with no detectable lesions. These results suggest that bacterial community structure is already significantly altered in the colons of ACF patients.

**Figure 2.**
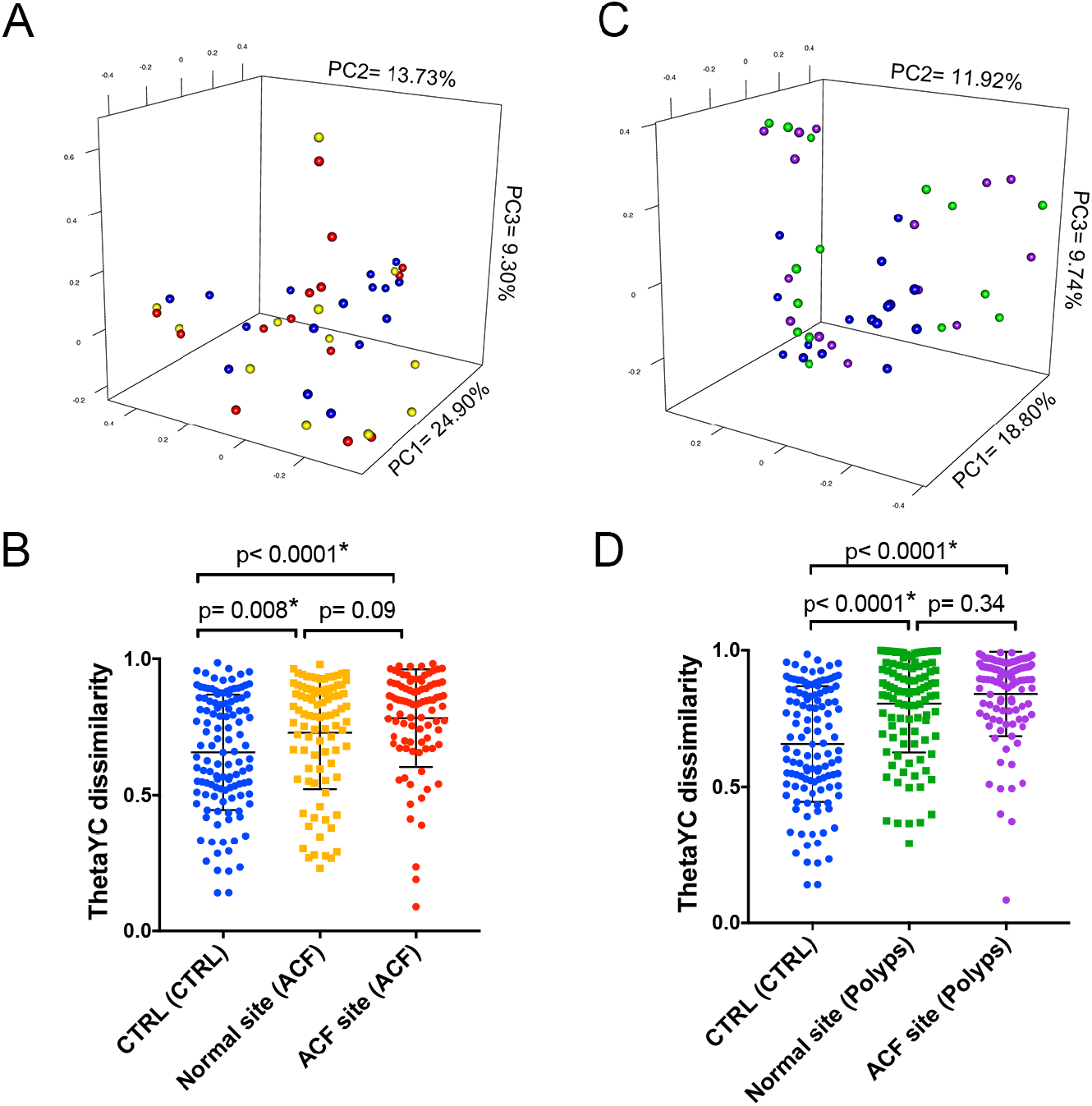
Beta diversity comparison of the microbiome between normal mucosa and ACF based on theta-YC distance. In polyp-free subjects, principle coordinate analysis (PCoA) plots showed no significant spatial separation as determined by AMOVA between ACF lesions and control mucosa taken from both ACF-free control and paired normal samples from the same ACF subject (A). The distance measured pair-wise between each sample, however, showed significant differences in microbiome community structure (B). In polyp subjects, PCoA plots showed no significant spatial separation tested using AMOVA between ACF lesions and control mucosa from both ACF-free controls and paired normal samples taken from the same ACF subject (C). However, spatial separation was more distant from control samples (blue) compared to A. In addition, the distance measured pair-wise between each sample showed significant differences in microbiome community structure (D), with more significant p-values from B when comparing the ACF site from polyp subjects (purple) to control group (blue) using Mann-Whitney test.

Next, in order to determine whether microbiome changes are associated with lesion progression, we analyzed a subset of ACF and their adjacent normal mucosal samples from patients with synchronous polyps. Interestingly, the PCoA plot showed a clearer separation of clustering of control mucosal samples (blue) from ACF samples (purple) or their adjacent normal mucosa (green) (p= 0.004) (**Figure 2C**). Microbiome alterations that were observed in ACF samples with synchronous polyps were greater than those changes found in ACF samples without synchronous polyps. The thetaYC dissimilarity comparison confirmed a greater difference between control normal samples and ACF samples with synchronous polyp(s) present (p< 0.0001) (**Figure 2D**), implicating further mucosal dysbiosis associated with lesion progression in the gut.

### ACF-adherent bacterial taxa are stratified by the presence of synchronous polyps and somatic mutations

We further identified differentially abundant specific bacterial taxa present within ACF lesions from patients with and without synchronous proximal polyps (**Figure S3**). The top 15 most differentially abundant taxa in ACF lesions taken from patients without synchronous proximal polyps included a significant increase of *Lactonifactor sp*. (OTU036) and *Pantoea sp*., (OTU038), as well as a significant decrease of *Clostridium XIVa sp*. (OTU103); however, each of the 15 taxa yielded a relative abundance of less than 1%. On the other hand, the 15 most differentially abundant taxa present in ACF lesions taken from patients with synchronous proximal polyps included a significant increase of *Eubacterium sp.* (OTU060) and *Clostiridium sensu stricto sp.* (OTU087). Notably, the levels of *Faecalibacterium sp.* (OTU001) were significantly lower in mucosal samples taken from patients without synchronous polyps compared to normal mucosa from control subjects.

Next, we examined the possibility that the presence of a somatic mutation in ACF may influence the association and composition of colon-associated bacterial species. To test this possibility, host DNA was examined by DNA mass-spectrometry analysis using our recently reported customized CRC somatic mutation panel ^9^. Patients harboring a somatic mutation to the tumor suppressor gene, *APC* (*R1450*, R876*, S1465fs\**3), showed a significant correlation with *Intestinibacter sp*. (p< 0.001). ACF with a mutation in the proto-oncogene, *KRAS* (G12V, G12D), showed a significant correlation with *Ruminococcus gnavus* (p< 0.001) (**Figure 3**). Although limited in sample size, these results provide the first indication of a potential relationship between specific adherent bacterial species and a particular CRC-related somatic mutation.

**Figure 3.**
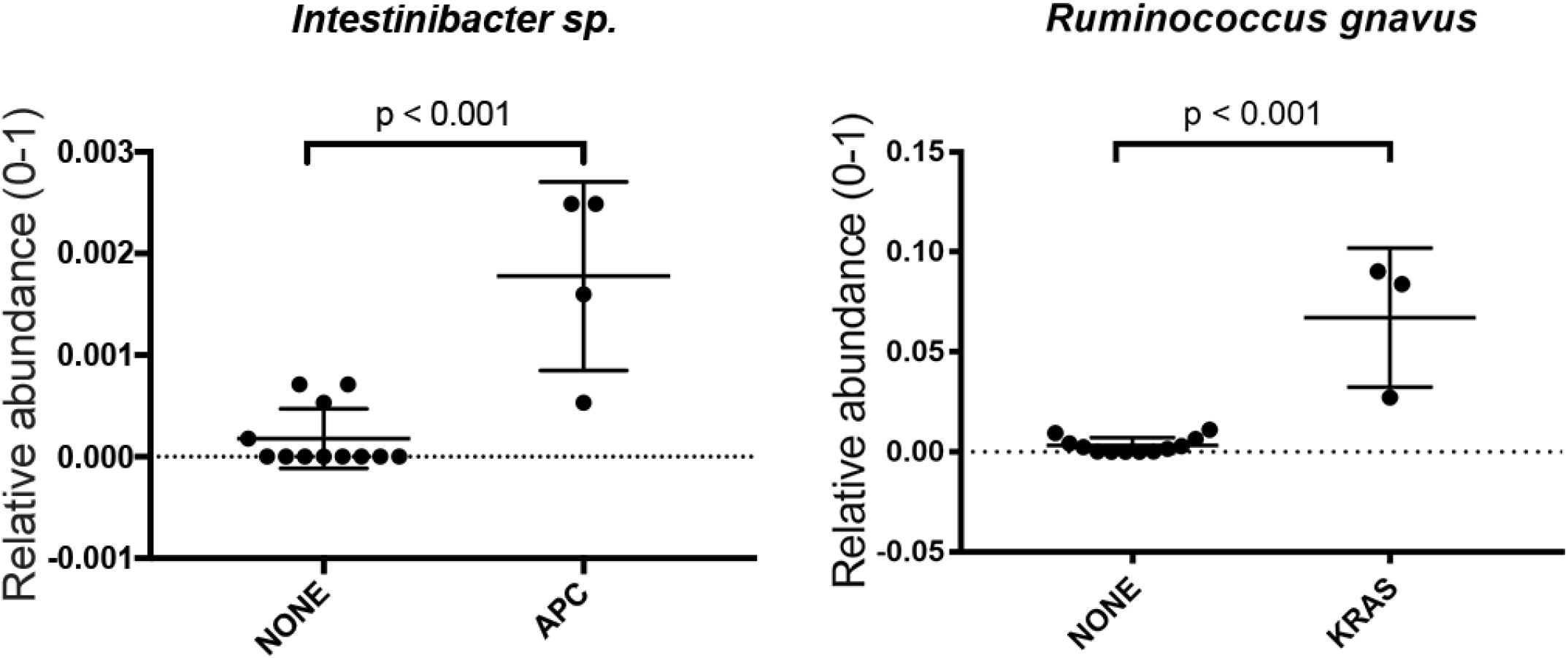
Significantly abundant bacterial taxa by DNA mutation types. *Intestinibacter sp.* was significantly increased in ACF samples with an *APC* mutation, while *Ruminococcus gnavus* was significantly increased in ACF samples with a *KRAS* mutation compared to samples with no mutation detected.

### Two distinct types of ACF bacterial communities (Microbiome Cluster A and B) and their correlation to clinical parameters

We further investigated the extent to which the microbiome community signature may shape the potential progression of ACF lesions into more advanced polyps (sessile serrated adenomas and polyps (SSA/P), hyperplastic polyps (HP), or traditional adenomas (TA)). **Figure 4** shows ACF bacterial profiles depicted as a heat-map, with the demographic and clinical characteristics of the patients denoted. Two distinct microbiome clusters segregated by bacterial community profile were identified in ACF lesions; 62 samples were present in Microbiome Cluster A and 12 samples were present in Microbiome Cluster B. The presence of an ACF lesion alone did not distinguish Microbiome Clusters A and B (p= 0.4089). However, an ACF lesion with or without the presence of a synchronous polyp clearly distinguished Microbiome Clusters A and B (p< 0.0001), indicating that a distinct bacterial community composition may play a role in the eventual progression of ACF to polyps. Importantly, more than 50% of ACF samples within Microbiome Cluster A occurred in the absence of synchronous polyps, whereas more than 90% of ACF samples within Microbiome Cluster B had at least one synchronous proximal polyp present in the same subject.

**Figure 4.**
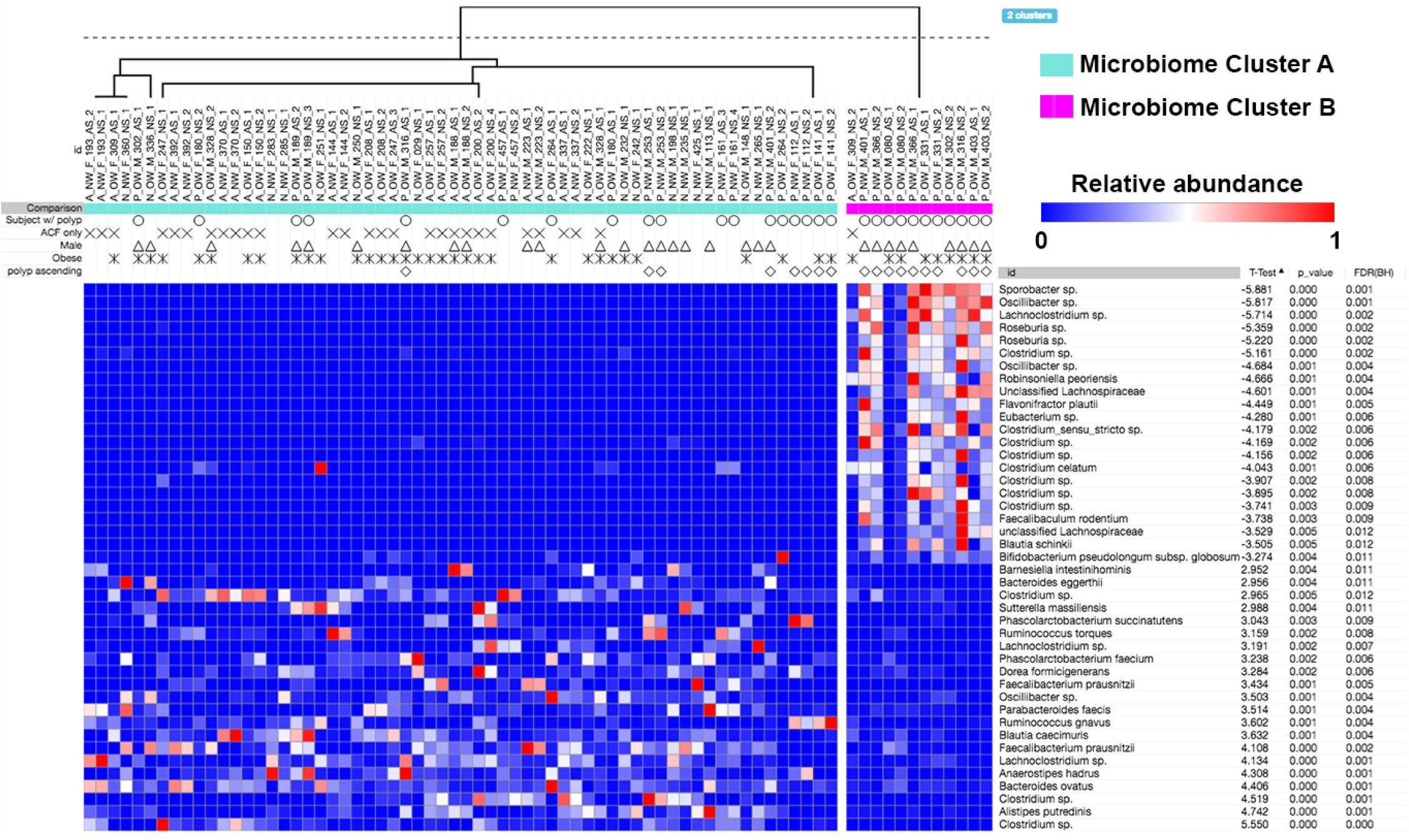
Microbiome profiles of the entire sample set. Heat-maps were generated using Morpheus based on the top 100 OTUs found in the biopsy specimens. Samples are shown in columns, while each OTU is depicted in the rows. Only significantly different taxa in each cluster are shown. The color scale appears on the top right side of the Figure. Unsupervised hierarchical clustering (complete linkage) shows two clusters, Microbiome Cluster A (cyan) and Microbiome Cluster B (magenta). Annotation for the presence of a polyp(s) in the subject, gender and obesity, as well as the presence of a polyp(s) in the ascending colon are indicated by unique shapes when positive. Significance was determined after a Benjamini-Hochberg multiple comparison adjustment.

Based on the strong association for the presence of a synchronous polyp with Microbiome Cluster B, we further investigated the location and subtype of polyps associated with this cluster (**Figure 5**). Out of the 12 samples represented within this group, 2 samples had sessile serrated adenomas, 7 samples had tubular adenomas and 2 samples had hyperplastic polyps, whereas the one remaining sample in this cluster had no evidence of a synchronous polyp. Overall, Microbiome Cluster B was significantly correlated with the presence of a polyp in the proximal colon (p= 0.0073). Furthermore, all normal mucosal samples were found within Microbiome Cluster A. Although our sample sizes are somewhat limited, we believe these results suggest that the Microbiome Cluster B profile may be associated with a mucosal dysbiosis that contributes to proximal colon lesion progression.

**Figure 5.**
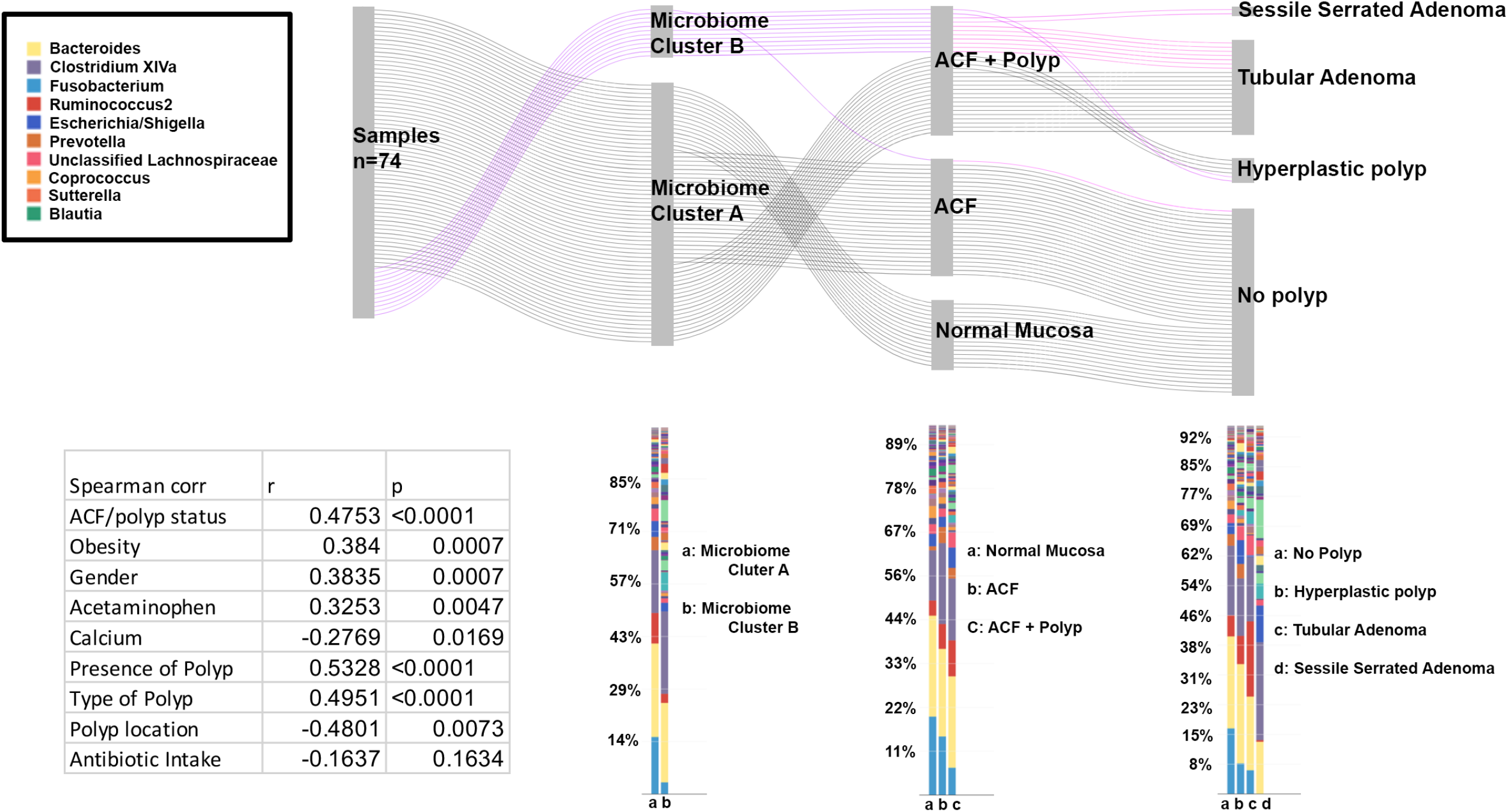
Microbiome signature shapes the development of polyps. Two distinct microbiome clusters (Microbiome Clusters A and B) defined by bacterial community signature were depicted in the Sankey diagram, demonstrating the ‘flow’ of each biopsy sample. Microbiome Cluster B is an active driver in the progression of pre-cancerous lesions. Over 90% of samples with Microbiome Cluster B yielded malignant polyps in the same subject. Less than 50% of samples with Microbiome Cluster A were associated with the presence of any polyp in the same subject (Top). Bacterial relative abundances in each category were depicted in the bar graph comparing groups from the Sankey diagram above. There were bacterial community differences by Microbiome Cluster type, ACF/polyp status (no lesion, ACF, and ACF with polyp), and the specific type of polyp present in the subjects. Microbiome Clusters A and B were significantly correlated with demographic or clinical characteristics, such as obesity status, gender, acetaminophen and calcium intake. Antibiotic intake was not significantly correlated with Microbiome Clusters A and B (**Table 1**).

As further shown in **Figure 5**, known CRC risk factors, including gender, obesity, and polyp location were significantly correlated with Microbiome Cluster B (p= 0.0007, p= 0.0007, p= 0.0073 for these three risk factors, respectively). These results suggest that the observed distinct microbiome clusters are in concordance with a combination of CRC risk factors. Interestingly, acetaminophen intake was positively correlated with Microbiome Cluster B (p= 0.0047), whereas calcium consumption was negatively correlated with Microbiome Cluster B (p= 0.0169). It is noteworthy that antibiotic intake did not significantly correlate with either Microbiome Cluster A or B (p= 0.163).

### Predictive metabolic function

Several functional KEGG pathways of potential clinical significance were predicted within bacterial communities present between Microbiome Clusters A or B (**Figure 6**). Microbiome Cluster A showed functional changes associated with lipid metabolism, amino acid metabolism, nucleic acid metabolism, carbohydrate metabolism and metabolism of co-factors and vitamins, as well as DNA replication, repair, folding, sorting and degradation, and translation. On the other hand, Microbiome Cluster B was associated with the following predicted pathways: cell motility, membrane transport and signal transduction.

**Figure 6.**
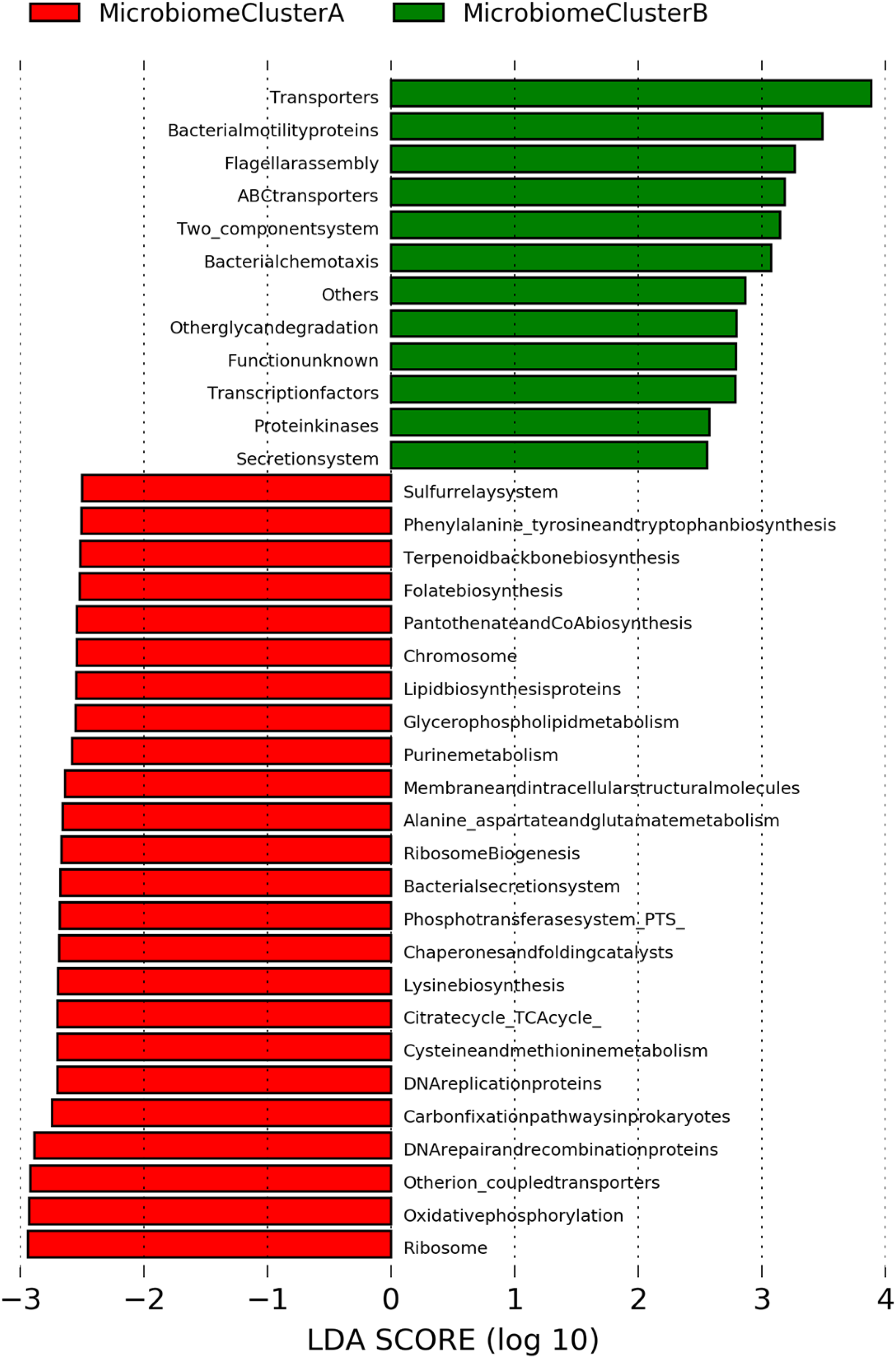
Predictive functional profiling annotation between Microbiome Clusters A and B. 16S rRNA gene sequencing data were clustered as OTUs with 97% identity, and predictive functional profiling was performed *via* PICRUSt using KEGG KO as a reference. Several virulence factors were predicted as functional pathways associated with Microbiome Cluster B (green).

## Discussion

In this study, we examined the potential influence of the microbiome on the pathogenesis of early colonic neoplasia. We determined the composition of the colon-associated microbiome in ACF and its relationship to the presence of synchronous polyps found in the proximal colons of normal colonoscopy screening subjects. Our long-term goal is to uncover novel microbiota-epithelial interactions, and further understand how microbial dysbiosis may impact the microenvironment of the colon. It is likely that specific microbiota-epithelial interactions may directly influence the growth potential of colonic ACF. The present study begins to develop a comprehensive data-set of precancerous lesions present in the proximal colon, including their histological features and associated somatic mutations that can be combined with detailed information on microbial community structure. We believe this combinatorial approach will enable a sensitive and comprehensive prediction of early cancer risk, and may ultimately provide new avenues for cancer prevention strategies based upon the targeted manipulation of the gut microbiota.

Previously, *F. nucleatum* was identified as one of the most abundant bacterial species associated with CRC tumors ^22,40–44^. Our data also indicate that ACF lesions display evidence of dysbiosis of colon-associated mucosal bacteria. However, the specific taxa shown to be associated with later stages of CRC, such as the aforementioned *F. nucleatum*, were neither prevalent nor abundant in our sample cohort. We believe our findings are consistent with those of a recent study ^45^ showing that *Fusobacterium nucleatum* stimulates growth of colorectal cancer cells without affecting precancerous adenoma cells, leading the authors to speculate that *F. nucleatum* may provide a ‘second hit’ to the initiated colonic epithelium. Regardless, our results clearly demonstrate significant dysbiosis of mucosal adherent microbiota, with an increase in presumed pathogens and a decrease in a number of taxa that may have beneficial properties. This latter group of microbiota may also include a significant decrease of *F. prausnitzii*, one of the more abundant taxa shown to have anti-inflammatory activity in the human intestine ^46,47^. Overall, our results suggest that dysbiosis of the colonic mucosal microbiome may contribute in part to the development of more advanced forms of colonic neoplasia.

Our results further demonstrate that it is possible to provide a detailed view of tightly associated microbial composition obtained from even the smallest mucosal biopsy specimens (~2-mm^3^), even after a thorough pre-colonoscopy purging protocol. Our sample set has readily provided several distinct microbial clusters, referred to as Microbiome Clusters A and B, showing a clear distinction among the three patient groups. Microbiome Cluster B has a particularly strong association with the presence of synchronous polyps in the proximal colon, including SSAs, TAs, and HPs. We further identified a group of strongly associated taxa with Microbiome Cluster A.

Based on *in silico* analysis, we are well aware of the relatively low taxonomic resolution resulting from the V4 hypervariable region of the 16S rRNA gene (data not shown). In fact, many of the OTUs identified by our analysis were classified at the genus level without inclusion of species names. However, the present study provides the first OTU-level bacterial community analysis associated with early neoplastic changes occurring within the human colonic mucosa.

Predictive metabolic function analysis demonstrated that the functional categories associated with Microbiome Cluster B uniquely included a group of virulence factors, including bacterial motility proteins, two-component system, secretion system, flagella assembly, and chemotaxis. These results are consistent with a previous study that demonstrated many similar predicted metabolic pathways in colorectal tumors ^48^. The metabolic function represented in Microbiome Cluster B may implicate bacterial community composition and their biological markers, and this may further explain the extent of mucosal dysbiosis uniquely observed in Microbiome Cluster B, even in the earliest pre-cancerous ACF lesions.

To gain further insight into how these microbial-epithelial interactions may impact early colonic neoplasia, we performed an analysis of CRC-related somatic mutations on a subset of laser-captured ACF samples using our customized CRC somatic mutation panel and DNA-Mass spectrometry ^9^. We found that *Intestinibacter sp*. and *Ruminococcus gnavus* are strongly associated with the presence of either an *APC* or *KRAS* mutation, respectively (**Fig. 3**). These results provide what we believe is the first direct evidence that cancer-related somatic mutations present at the earliest stages of colonic neoplasia may be directly associated with the presence of specific microbial organisms. Although these findings are suggestive, the possibility of a direct host-microbial interaction remains unknown. We believe these findings certainly warrant further investigation into potential mechanisms through which host-microbial interactions may contribute to neoplastic progression.

In summary, our study provides the first comprehensive characterization of a distinct microbiome profile directly associated with ACF, the earliest morphologically identifiable lesion in the colon that likely precedes the formation of more advanced forms of colonic neoplasia. We believe that our emphasis on the characteristics of the microbiome during early neoplasia will have a significant impact in terms of clinical application. Capturing integrated and combined characteristics of pre-cancerous lesions, ranging from somatic mutations, histology and microbiome composition, will eventually lead to a more sensitive and comprehensive detection of cancer risk, and contribute to efforts in cancer prevention and advanced colonoscopy screening based on the gut microbiome.

## Acknowledgements

We thank Microbial Analysis and Service Core at The University of Connecticut, Storrs for 16S rRNA sequencing and Dr. Prashant Singh from the Roswell Park Comprehensive Cancer Institute for the somatic mutation analysis using DNA MassArray. This work was supported by The State of Connecticut Department of Public Health, Biomedical Trust Fund (DWR), The American Institute for Cancer Research (DWR) and Institutional support from The Jackson Laboratory of Genomic Medicine (GW).

## Authors’ contributions

HB, GMW, and DWR designed the study. JB and TD performed the colonoscopies and collected biopsy specimens. FF confirmed the histology of the biopsies. TI, YI, YT and AM performed the experiments. HB, TI, YT, MD, GMW, and DWR analyzed, interpreted the data and contributed to the preparation of the manuscript. All authors read and approved of the final version of the manuscript.

## LEGENDS TO SUPPLEMENTARY FIGURES

**Figure S1. Direct visualization of colon-associated bacteria present within the colonic mucosa of representative polyp tissues.** Synchronous polyps obtained from patients in Group III were directly examined for the presence of adherent bacteria using 16S universal FISH probes. Red: bacteria (EUB338-cy3 probe), Magenta: E-cadherin, Green; Mucin-2, Blue: DAPI. Bacteria are depicted by white arrows. Bacteria were observed within the mucous layer associated with the epithelium.

**Figure S2. Study design and bacterial diversity comparison by biopsy location.** (A) Normal mucosal samples were taken from ACF-free and polyp-free subjects (blue). ACF lesions were biopsied from polyp-free subjects (red), as well as subjects with proximal polyp(s) present (purple). (B) Paired normal mucosa was biopsied from polyp-free subjects (yellow) and subjects with polyp(s) present in the proximal colon (green) in order to compare the microbiome of different sites within the same subject. No statistical differences were found across all sample sites using measurements of bacterial richness (number of taxa found), evenness and diversity based on Shannon diversity and evenness indices in any pair of comparison groups.

**Figure S3. Differentially abundant OTUs between ACF, with and without synchronous polyps.** The top 15 differentially abundant OTUs occurring between ACF biopsies without a polyp (left panel) and with a synchronous polyp (right panel) compared to normal mucosa taken from the control subjects. Individual taxa were plotted separately (bottom) for taxa showing significance between ACF and normal mucosa from control subjects using Wilcoxon-signed rank test. *Indicates a p-value less than 0.05, **indicates a p-value less than 0.005

**Figure S4.** Top 3 most abundant phyla between Microbiome Clusters A and B were characterized at the genus taxonomic level. Firmicutes, Bacteroidetes and Proteobacteria were the most abundant phyla in both cluster types. The most distinct bacterial composition observed between two clusters was within the Firmicutes phylum. Microbiome Cluster A showed *Faecalibacterium* as a dominant genus, while Microbiome Cluster B showed *Clostridium XIVa* as a dominant genus.

