## Supplementary figures and images for "Characterization of Mucosal Dysbiosis of Early Colonic Neoplasia"

### SF1

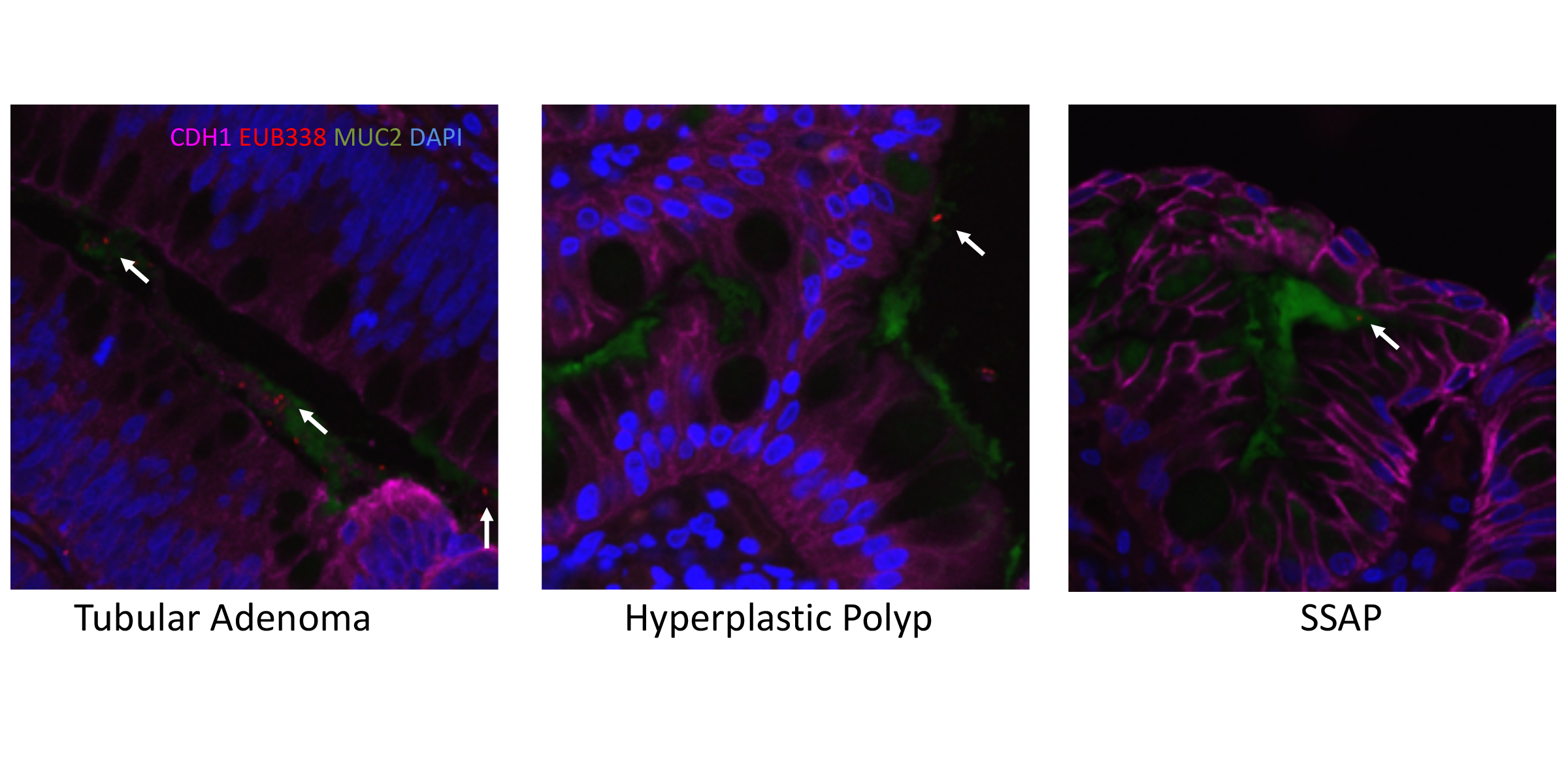

### SF2

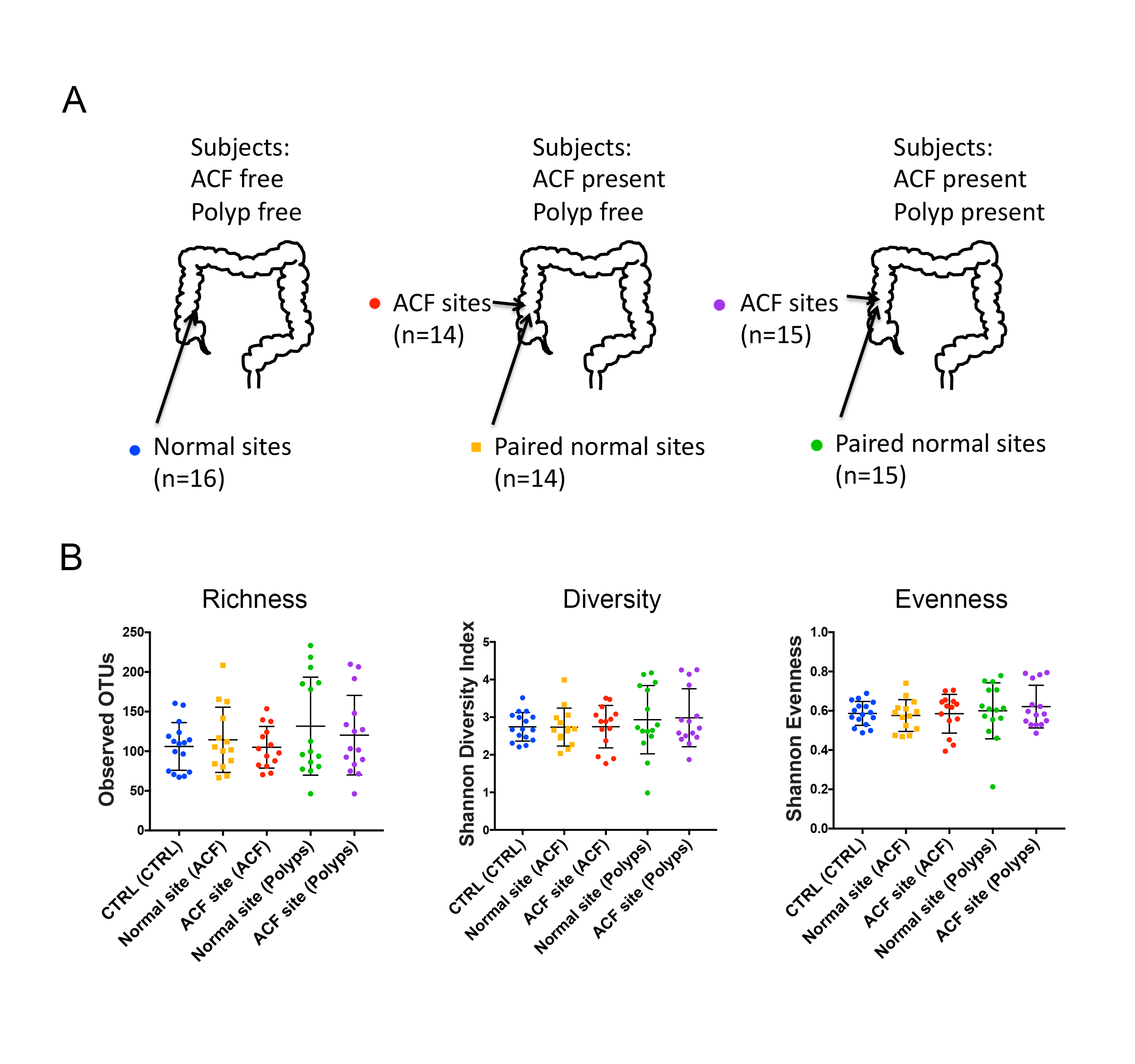

### SF3

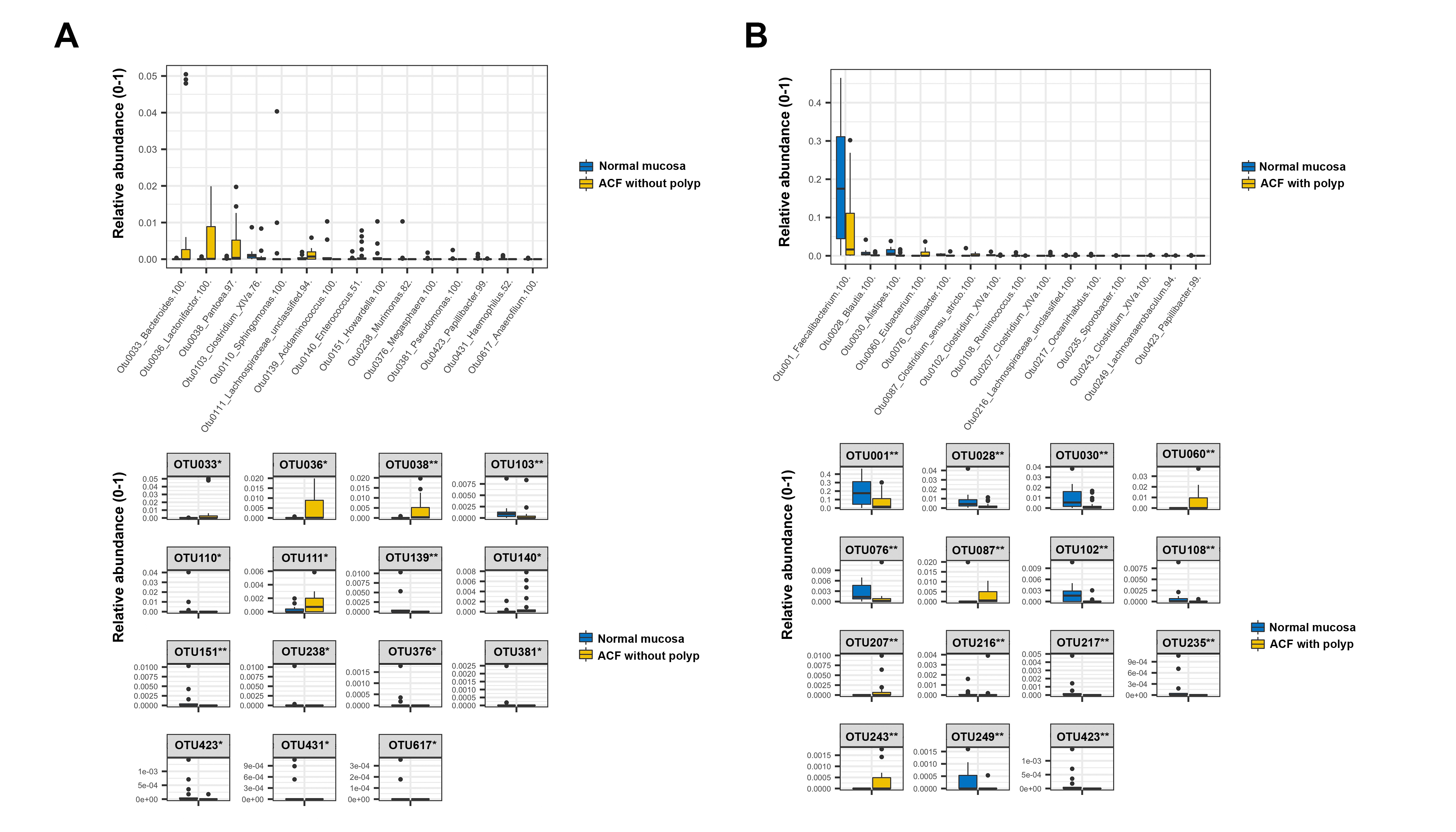

### SF4

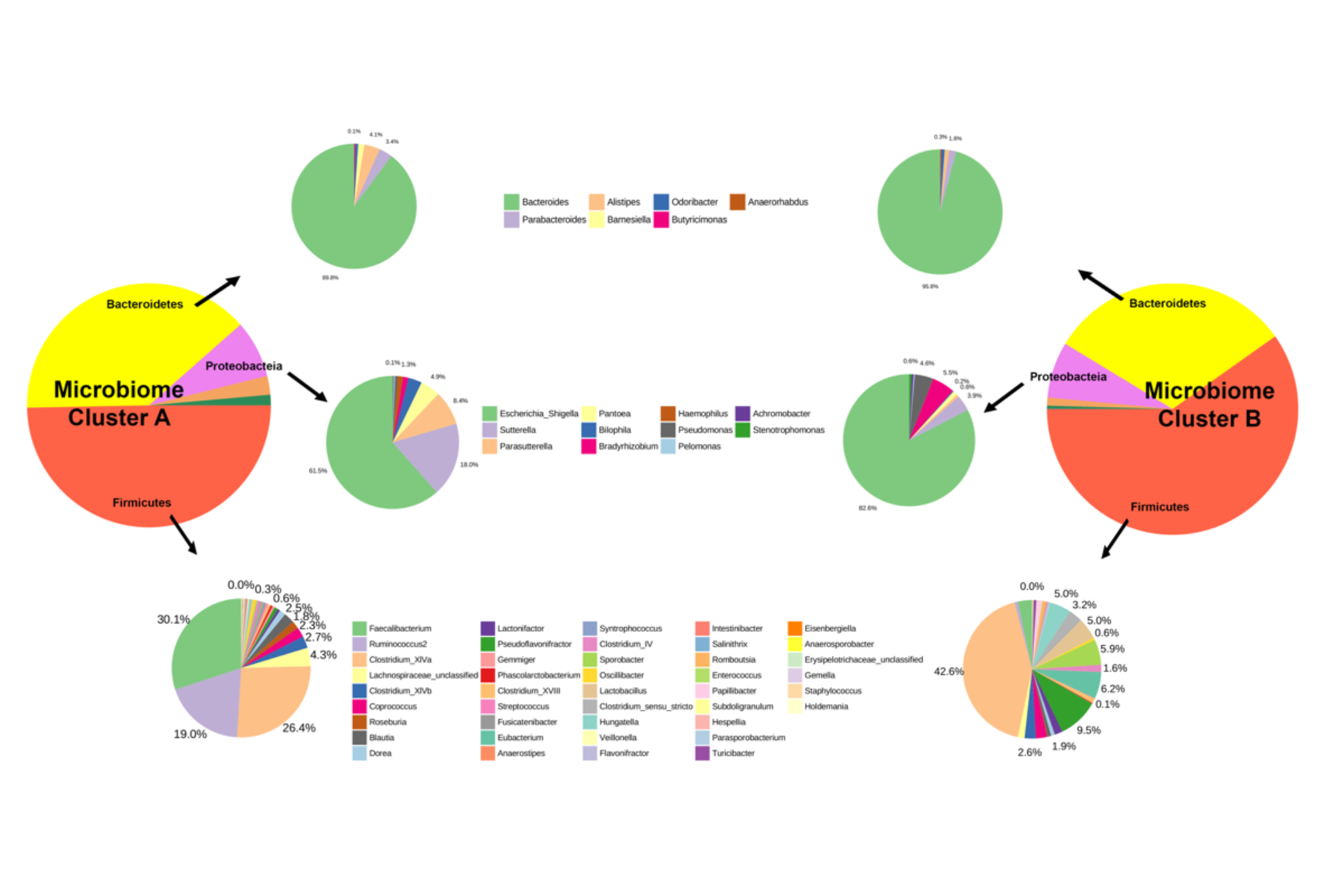
